# Small-molecule ligands can inhibit −1 programmed ribosomal frameshifting in a broad spectrum of coronaviruses

**DOI:** 10.1101/2021.08.06.455424

**Authors:** Sneha Munshi, Krishna Neupane, Sandaru M. Ileperuma, Matthew T.J. Halma, Jamie A. Kelly, Clarissa F. Halpern, Jonathan D. Dinman, Sarah Loerch, Michael T. Woodside

## Abstract

Recurrent outbreaks of novel zoonotic coronavirus (CoV) diseases since 2000 have highlighted the importance of developing therapeutics with broad-spectrum activity against CoVs. Because all CoVs use −1 programmed ribosomal frameshifting (−1 PRF) to control expression of key viral proteins, the frameshift signal in viral mRNA that stimulates −1 PRF provides a promising potential target for such therapeutics. To test the viability of this strategy, we explored a group of 6 small-molecule ligands, evaluating their activity against the frameshift signals from a panel of representative bat CoVs—the most likely source of future zoonoses—as well as SARS-CoV-2 and MERS-CoV. We found that whereas some ligands had notable activity against only a few of the frameshift signals, the serine protease inhibitor nafamostat suppressed −1 PRF significantly in several of them, while having limited to no effect on −1 PRF caused by frameshift signals from other viruses used as negative controls. These results suggest it is possible to find small-molecule ligands that inhibit −1 PRF specifically in a broad spectrum of CoVs, establishing the frameshift signal as a viable target for developing pan-coronaviral therapeutics.

---

The 21^st^ century has seen a series of public-health emergencies caused by zoonotic coronavirus (CoV) diseases: the SARS epidemic in 2002–3, periodic MERS outbreaks since 2012, and the ongoing COVID-19 pandemic (1). Given ever-increasing human contact with major CoV reservoirs like bats (2) due to habitat encroachment and climate change, novel CoV diseases will likely continue to emerge in the near future, generating new public-health challenges. It is therefore urgent to find anti-viral therapeutics that are effective against a broad spectrum of CoVs, especially bat CoVs, from which the previous 21^st^-century CoV zoonoses are thought to have originated and which are one of the most likely sources for future novel CoVs (2,3). To date, however, no drugs proven to be effective against a broad spectrum of CoVs have been identified.

One promising target for developing broad-spectrum CoV therapeutics is a process that plays a key role in gene expression in all CoVs: −1 programmed ribosomal frameshifting (−1 PRF). The proteins needed for transcription and replication of the viral RNA in CoVs are encoded in ORF1b, which is out of frame with respect to ORF1a, and they are only expressed if the ribosome shifts into the −1 frame at a specific location in the viral genome (4). This reading-frame shift is directed by a frameshift signal in the mRNA that consists of a ‘slippery sequence’ where the reading-frame shift occurs and a down-stream structure in the mRNA that stimulates the frameshift, separated by a ∼5–7-nucleotide (nt) spacer (5-7). Altering the level of −1 PRF by mutations or ligands that bind to the stimulatory structure has been shown to attenuate the virus significantly for SARS-CoV and SARS-CoV-2 (8-11), as well as for other viruses dependent on −1 PRF (12-14), leading to efforts to find ligands with potential therapeutic activity.

In the case of CoVs, the stimulatory structure is an RNA pseudoknot, a structure formed from 2 or more interleaved hairpins in which the loop of one hairpin is base-paired to form the stem of another (Fig. 1A). Most drugs target proteins, because of their well-defined structures and binding pockets allowing for high-affinity and high-specificity binding. Because RNA structures like pseudoknots also offer complex surfaces with binding pockets for potential ligands, they are also well-suited for druggability (15-17). Indeed, CoV pseudoknots feature an unusual 3-stem architecture that is more complex than the 2-stem architecture typical of most stimulatory pseudoknots (18), as illustrated in recent studies of the SARS-CoV-2 pseudoknot (Fig. 1B) (10,19-21); this complexity creates a number of putative binding pockets (22) and reduces the likelihood of off-target binding. Moreover, the pseudoknot sequence is generally highly conserved in CoVs (23), suggesting not only that many CoV pseudoknots may share structural features that could allow for ligands to bind a broad spectrum of CoV pseudo-knots, but also that they may be less susceptible to mutations that induce drug resistance by altering the pseudoknot structure. As an additional benefit, targeting the −1 PRF signal is orthogonal and complementary to more standard strategies of targeting viral proteins such as the RNA-dependent RNA polymerase (RdRP) or viral proteases, holding out the promise for effective combination therapies.

**Figure 1:**
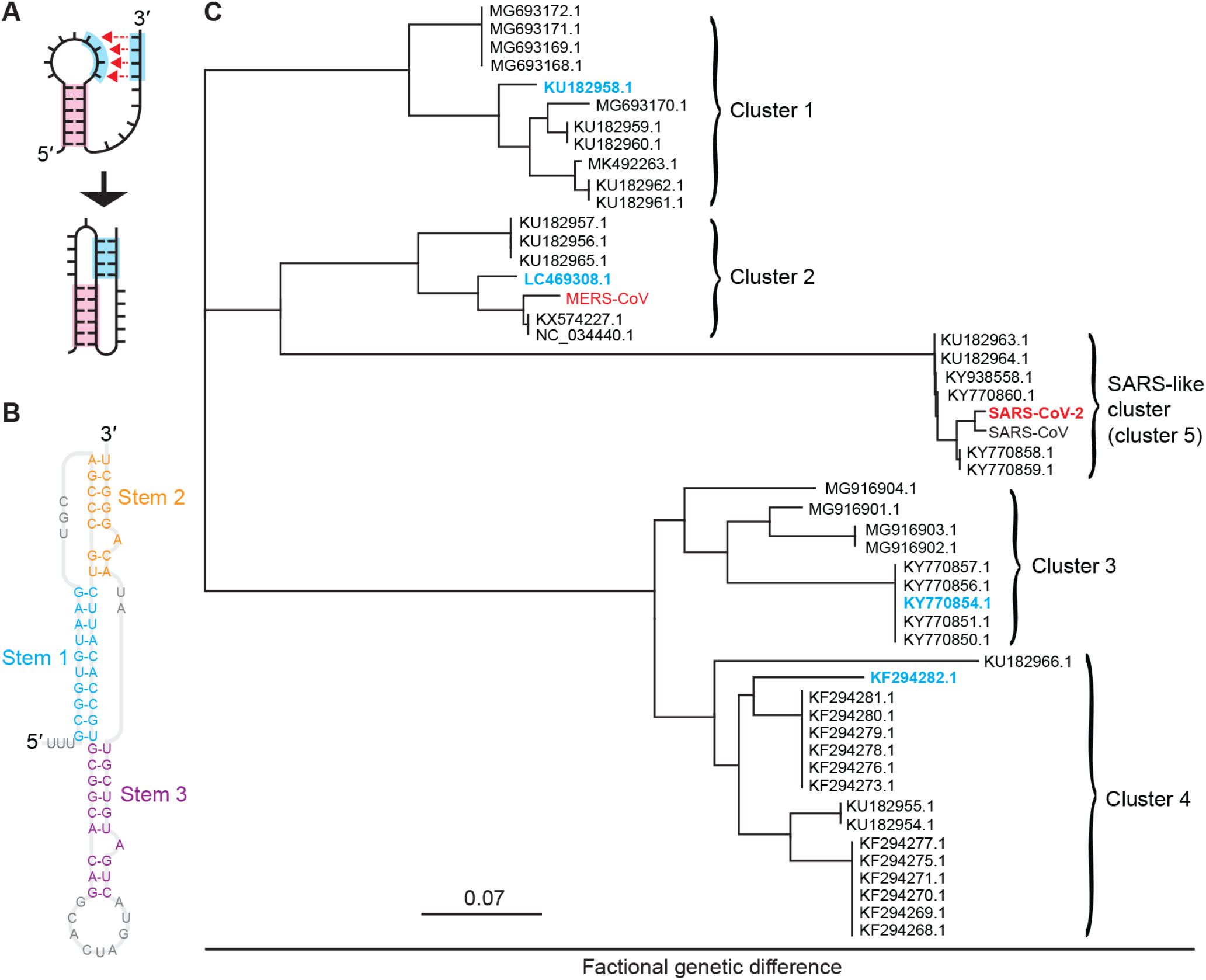
Pseudoknots from bat coronaviruses. (A) Pseudoknots form when the unpaired bases in an RNA stem loop pair with another single-stranded segment. (B) CoV pseudoknots have a 3-stem architecture, illustrated here for the pseudoknot from SARS-CoV-2. (C) Phylogenetic tree showing five clusters of bat and human CoVs with similar pseudoknot sequences. Representatives from each cluster studied here are shown in cyan for bat CoVs and red for human CoVs.

Several studies have identified small-molecule ligands that modulate −1 PRF in human CoVs. Computational docking against the SARS-CoV pseudoknot found a compound, 2-[{4-(2-methyl-thiazol-4ylmethyl)-[1,4]diazepane-1-carbonyl]-amino}-benzoic acid ethyl ester (hereafter denoted MTDB) that inhibited −1 PRF in SARS-CoV (24,25), SARS-CoV-2 (19), and mutant variants of SARS-CoV-2 (26), and also suppressed viral replication (10). More recently, empirical screens for modulation of −1 PRF in SARS-CoV-2 have found a number of compounds that either enhance or suppress −1 PRF (11,27,28), including merafloxacin, a fluoroquinolone antibacterial that was also effective at suppressing viral replication and showed resistance to natural mutants of the pseudoknot (11). Anti-sense oligomers have also been explored for modulating −1 PRF in SARS-CoV (29), MERS-CoV (30), and SARS-CoV-2 (21). However, little has been reported about frameshifting in bat CoVs, which are the likely source of 5 of the 7 existing human CoVs. Although many bat-CoV sequences have been determined and analyzed to identify putative −1 PRF signals (23), frameshifting by these −1 PRF signals has not yet to our knowledge been confirmed experimentally. Furthermore, no pseudoknot structures from bat CoVs have been reported yet, nor have any studies of inhibitors targeting frameshifting in bat CoVs.

Here we examine −1 PRF in bat CoVs, comparing it to −1 PRF in bat-derived human CoVs like SARS-CoV-2 and MERS-CoV. Using dual-luciferase assays of frameshifting, we first confirmed the activity of the putative −1 PRF signals from a panel of four CoVs chosen to be representative of the range of sequences found to date across bat CoVs. We then augmented the pool of potential −1 PRF inhibitors to use in the study by screening FDA-approved drugs for activity against −1 PRF in SARS-CoV-2. Choosing six small-molecule ligands confirmed by us and others to inhibit −1 PRF in SARS-CoV-2, we compared their effects on −1 PRF induced by a panel of six CoV frameshift signals, from four bat CoVs as well as MERS-CoV and SARS-CoV-2. We observed that −1 PRF could be inhibited substantially for each of the bat-CoV frameshift signals, suggesting that they can all be targeted therapeutically. Moreover, several of these compounds had moderate to strong activity against more than one CoV; one, nafamostat, was active to some degree against all of the CoV frameshift signals tested. These results suggest that CoV frameshift signals are a viable target for developing novel broad-spectrum anti-CoV therapeutics.

## METHODS

### Clustering of bat coronavirus genomes

All known bat coronavirus sequences were downloaded on April 30, 2020 as an alignment from the NCBI Virus database. These were imported into Geneious Prime version 2020.1.2 (www.geneious.com), where the frameshift region was extracted. Only sequences with coverage in the frameshift region were retained for clustering. A phylogenetic tree was created based on intersequence distances at the frameshift site, revealing five clusters (Fig 1C). Secondary structural predictions were performed on the pKiss (31) and Hotknots (32,33) web servers using default settings, with the proposed structures (Fig. S1) selected as the lowest-energy consensus results homologous to the SARS-CoV and SARS-CoV-2 pseudoknots.

### Preparation of mRNA constructs

(1) Constructs for testing inhibitors. For measuring −1 PRF efficiency in the panel of CoV frameshift signals used to test inhibitors, we used a dual-luciferase reporting system based on a plasmid containing the sequence for *Renilla* luciferase and the multiple cloning site (MCS) from the plasmid pMLuc-1 (Novagen) upstream of the firefly luciferase sequence in the plasmid pISO (addgene), as described previously (19,26). The frameshift signals for all CoVs in the testing panel as well as the two viruses used as negative controls (HIV-1 and PEMV1) were cloned into the MCS between the restriction sites PstI and SpeI. Three different types of constructs were made for each CoV frameshift signal. First, a construct for assaying −1 PRF efficiency was made: it contained the frameshift signal with slippery sequence (UUUAAAC) and pseudoknot, and the downstream firefly luciferase gene was placed in the −1 frame so that its expression was dependent on −1 PRF. Next, we made two controls from this construct: (i) a negative control to measure the background firefly luciferase luminescence (0% firefly luciferase read-through), where the slippery sequence was mutated to include a stop codon (UUGAAAC); and (ii) a positive control to measure 100% firefly luciferase read-through, where the slippery sequence was disrupted (UAGAAAC) and the firefly luciferase gene was shifted into the 0 frame. Sequences of frameshift signals for all constructs are listed in Table S1. We amplified transcription templates from these plasmids by PCR, using a forward primer that included the T7 polymerase sequence as a 5′ extension to the primer sequence(19,26). We then produced the mRNA for dual-luciferase measurements from the transcription templates by *in-vitro* transcription (MEGAclear).

(2) Constructs for library screening. The dual-luciferase reporter system used for screening the drug library was based on previously described plasmids (19). Briefly, the frameshifting element from SARS-CoV-2—encompassing the attenuator hairpin, slippery sequence, spacer, and pseudoknot—was placed between the sequences for *Renilla* and firefly luciferases. For the construct used to measure −1 PRF, *Renilla* luciferase was in the 0 frame and firefly luciferase in the −1 frame. To measure 100% read-through, the firefly luciferase gene was placed in the 0 frame and the slippery sequence was disrupted by mutation as above. For the negative control, equivalent to 0% −1 PRF, we introduced silent mutations to the slippery sequence (CCUCAAC) that left the encoded protein sequence unchanged. Transcription templates were amplified from the plasmids by PCR and and transcribed *in vitro* with the HiScribe T7 High Yield RNA synthesis kit (New England Biolabs Inc.). To obtain sufficient amounts of mRNA, we scaled up our reactions five-fold compared to the manufacturer’s protocol. To remove rNTPs and polymerase, mRNA was purified using weak anion exchange as described (34).

(3) Construction of bi-fluorescent reporter for cell-based assays: The reporter plasmid pJD2261 was constructed by cloning a gBlock (IDT) encoding AcGFP, the HIV-1 −1 PRF signal, mCherry, and insulator A2 described previously(35) into PstI-digested pUC19 (Genbank accession L09137). Next, the CMV promoter from pJD2044 was inserted into KpnI/PstI-digested fluorescent reporter. The 0-frame control reporter was made by digesting pJD2261 with SalI and BamHI to remove the HIV-1 frameshift insert, gel-purifying the result, then ligating a DNA oligonucleotide insert (IDT) containing an α-helix spacer(36), with modified ends containing SalI and BamHI restriction sites. The SARS-CoV-2 frameshift reporter was made by digesting pJD2514 [19] using SalI and BamHI, gel-purifying the SARS-CoV-2 −1 PRF insert, and then ligating it into SalI/BamHI-digested pJD2261 vector. All reporter plasmids were sequence-verified. Oligonucleotide sequences for cloning are listed in Table S2.

### Dual-luciferase frameshift measurements

(1) Panel of CoV frameshift signals. The −1 PRF efficiency was measured for each member of the panel using a cell-free dual-luciferase assay (37). Briefly, for each construct, 1.2 µg of mRNA transcript was heated to 65 °C for 3 min and then incubated on ice for 2 min. The mRNA was added to a solution mixture containing amino acids (10 µM Leu and Met, 20 µM all other amino acids), 17.5 µL of nuclease-treated rabbit reticulocyte lysate (Promega), 5 U RNase inhibitor (Invitrogen), and brought up to a reaction volume of 25 µL with water. The reaction mixture was incubated for 90 min at 30 °C. Luciferase luminescence was measured using a microplate reader (Turner Biosystems). First, we mixed 20 µL of the reaction mixture with 100 µL of Dual-Glo Luciferase reagent (Promega) and incubated for 10 min before reading firefly luminescence, then we added 100 µL of Dual-Glo Stop and Glo reagent (Promega) to the mixture to quench firefly luminescence before reading *Renilla* luminescence. The −1 PRF efficiency was calculated from the ratio of firefly to *Renilla* luminescence, *F*:*R*, after first subtracting the background *F*:*R* measured from the negative control and then normalizing by the 100% readthrough *F*:*R* from the positive control.

To quantify the effects of the inhibitors on −1 PRF efficiency, compounds were added to the reaction volume at a final concentration of 20 μM for each construct (frameshift signals from CoVs, HIV, and PEMV1, as well as all positive and negative controls). Compounds were dissolved in DMSO, leading to a final DMSO concentration in the assays of 1% by volume. The results were averaged from 3–9 replicates, as described previously (19,26). The *Renilla* and firefly luciferase luminescence levels were similar for the different frameshift signals. The compounds did not affect the luminescence levels for *Renilla* luciferase, indicating that they did not interfere with translation, but rather their effects were specific to −1 PRF.

(2) Drug screen. The dual-luciferase assay in rabbit reticulocyte lysate (RRL) was optimized for a 384-well screening format by varying RRL content and salt concentrations, leading to conditions (reaction volume of 5 μL containing 20% RRL, with final concentrations of 25 mM potassium chloride, 0.5 mM magnesium acetate, and 65 mM potassium acetate, as well as 1.6 ng/ul mRNA, supplemented with 1 U/μl RNase inhibitor (RNaseIn, Promega), 20 μM amino acid mix (Promega), and 2mM DL-dithiothreitol) that yielded −1 PRF efficiencies comparable to the standard protocol supplied by the manufacturer (Fig. S1). For screening, a commercial library of 1,814 FDA-approved drugs (MedChem Express, Cat. No. HY-L022M) was reformatted into 384-well plates as 50 μM stock solutions in nuclease free-water with or without 1% DMSO, depending on the drug solubility. 1.0 μL of each compound was added to 4 μL of master mix containing all other reagents in 384-well flat white plates (Corning) on ice, for a final drug concentration of 10 μM. Positive and negative controls, as well as the frameshifting construct for normalization, were included in alternating wells of columns 1, 2, 23, and 24 on each plate, all treated with 1.0 μL of 1% DMSO. Plates were sealed, briefly mixed and centrifuged, and then incubated at 30 °C for 75 min. Reactions were quenched after incubation with 20 μM puromycin, and 20 μL of each luciferase substrate from the Luc-Pair Duo-Luciferase HT Assay kit (GeneCopeia) was added sequentially following manufacturer instructions. Luminescence was measured with 2 sec integration times in a microplate reader (Tekan Spark), 20 min after adding each substrate. The addition of reagents to the microplates was timed at each step to match the reading time delays and reading sequence of the microplate reader.

Z′-factors were determined for each plate using positive, negative and −1 PRF controls as previously described (38). The normalized −1 PRF inhibition was calculated by setting the average signal of the negative control (slippery-site mutant) to 100% inhibition, the average signal of the positive control (0 frame read-through) to 0%, and normalizing the result for each drug to the drug-free −1 PRF control.

All data analysis was carried out in Graphpad Prizm 8 and 9. We also tested for compounds that selectively inhibit firefly luciferase, which will lead to false positives, by incubating reactions without any drug present, quenching the reaction with puromycin before adding the compounds, and finally continuing as in the standard protocol.

### Bi-fluorescence frameshift measurements

A549 (CRL-185) cells were purchased from the American Type Culture Collection (Manassas, VA). Cells were maintained in F-12K media (Corning) supplemented with 10% fetal bovine serum (Gibco) and 1% penicillin-streptomycin and grown at 37°C in 5% CO_2_. Cells were seeded at 1×10^5^ cells per well in 12-well plates. Cells were transfected 24 hr after seeding with 0.75 µg of the bi-fluorescent reporter plasmid using Lipofectamine 3000 (Invitrogen) as per manufacturer’s protocol. Cells were treated 24 hr after transfection with a final concentration of 10 µM of the designated inhibitor, and then incubated for an additional 24 hours. Cells were collected by scraping into phosphate-buffered saline, pelleted by centrifugation, then lysed in Triton lysis buffer (1% Triton, 150 mM NaCl, 50 mM Tris pH 8, 1× Halt protease-inhibitor cocktail (Thermo Scientific)). Cell lysates were clarified by centrifugation and then assayed in a clear-bottom black-walled 96-well plate (Grenier Bio-One), quantifying fluorescence using a GloMax microplate luminometer (Promega). Fluorescence levels were measured using the “green” optical kit for mCherry (excitation at 525 nm, emission at 580–640 nm) and the “blue” optical kit for AcGFP (excitation at 490 nm, emission at 510– 570 nm). The −1 PRF efficiencies were calculated (after correcting for AcGFP fluorescence bleed-over into the mCherry channel) from the ratio of mCherry to AcGFP fluorescence, *mCh*:*AcG*, by first subtracting the background *mCh*:*AcG* measured from mock transfected cells and then normalizing by the 100% read-through *mCh*:*AcG* from the positive control.

## RESULTS

To build a panel of representative bat-CoV pseudoknots, we performed a multiple sequence alignment of 959 bat-CoV sequences from the NCBI virus database (39). Of these, 48 had significant coverage of the frameshift region. Calculating pairwise genetic distances between the frameshift signals, we used a multidimensional scaling algorithm (40) to collapse the genetic distances into a 2-dimensional display and thereby identified five distinct clusters. These clusters are illustrated in Fig. 1C: one (cluster 5) is closely related to SARS-CoV and SARS-CoV-2, and another (cluster 2) is closely related to MERS-CoV, whereas the other three clusters are either more distantly related or unrelated to human CoVs. For our testing panel, we chose one representative sequence from clusters 1 through 4 (Fig. 1C, cyan), as well as SARS-CoV-2 from cluster 5 (SARS-like cluster) and MERS-CoV from cluster 2 (Fig. 1C, red), in order to sample the breadth of sequence diversity and hence concomitant structural diversity among the frameshift signals. The choices for bat CoVs, denoted by their NCBI Virus accession numbers, were: KU182958, a beta-CoV isolated from the fruit bat *Rousettus leschenaultii* (cluster 1); LC469308, a beta-CoV isolated from the vesper bat *Vespertilio sinensis* (cluster 2); KY770854, an alpha-CoV isolated from the horseshoe bat *Rhinolophus macrotis* (cluster 3); and KF294282, an alpha-CoV isolated from the bent-wing bat *Miniopterus schreibersii* (cluster 4). The sequences used for each of the frameshift signals are listed in Table S1.

We first measured the basal efficiency of −1 PRF stimulated by each of the 6 frameshift signals from the testing panel using cell-free translation of a dual-luciferase reporter system (26,37), to confirm that they did indeed induce −1 PRF and assess the efficiency with which they did so. The reporter system consisted of the *Renilla* luciferase gene in the 0 frame upstream of the firefly luciferase gene in the −1 frame, with the two separated by the CoV frameshift signal in the 0 frame. The −1 PRF efficiency was obtained from the ratio of luminescence emitted by the two enzymes, as compared to controls with 100% and 0% firefly luciferase read-through. We found that every putative frameshift signal did indeed stimulate −1 PRF, but the efficiency varied widely: the two bat beta-CoV frameshift signals stimulated −1 PRF with ∼40% efficiency, comparable to SARS-CoV-2 and MERS-CoV, but the two bat alpha-CoVs did so at noticeably lower levels of ∼9% and 25% (Fig. 2).

**Figure 2:**
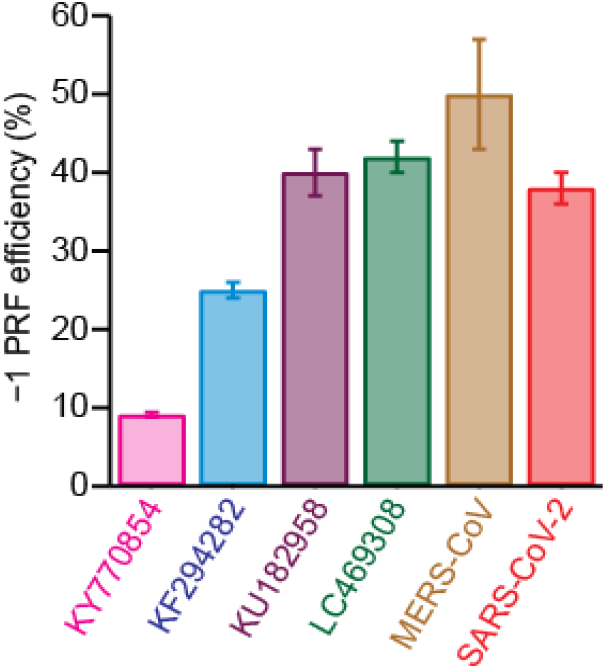
Efficiency of −1 PRF stimulated by bat CoV frameshift signals. The −1 PRF efficiency measured from cell-free dual-luciferase assays is in the range 25–50% typical of CoVs for all except KY770854 (cluster 3). Error bars represent standard error of the mean from 4–9 replicates.

We next investigated the effects of putative small-molecule inhibitors on −1 PRF efficiency for each frameshift signal in the panel. To identify additional inhibitors beyond those reported previously in the literature, we screened a library of 1,814 FDA-approved drugs to test their ability to modulate −1 PRF in SARS-CoV-2, using a cell-free dual-luciferase assay optimized for high-throughput screening (Fig. S2). Z′ factors of 0.67–0.82 for each plate suggested high quality and reproducibility of the assay (38). Selecting only those compounds that modulated −1 PRF by at least 30% or more, the initial screen identified 24 hits that had little to no effect on general protein translation (less than two-fold), as indicated by changes in the *Renilla* luminescence levels, and another 18 that also altered general translation levels (Table S3). After removing false positives that selectively inhibited firefly luciferase, we further validated hits from the initial screen in triplicate, confirming at least a 25% change in −1 PRF for eight of the compounds (Fig. 3A). Six of these inhibited −1 PRF, whereas two enhanced it.

**Figure 3:**
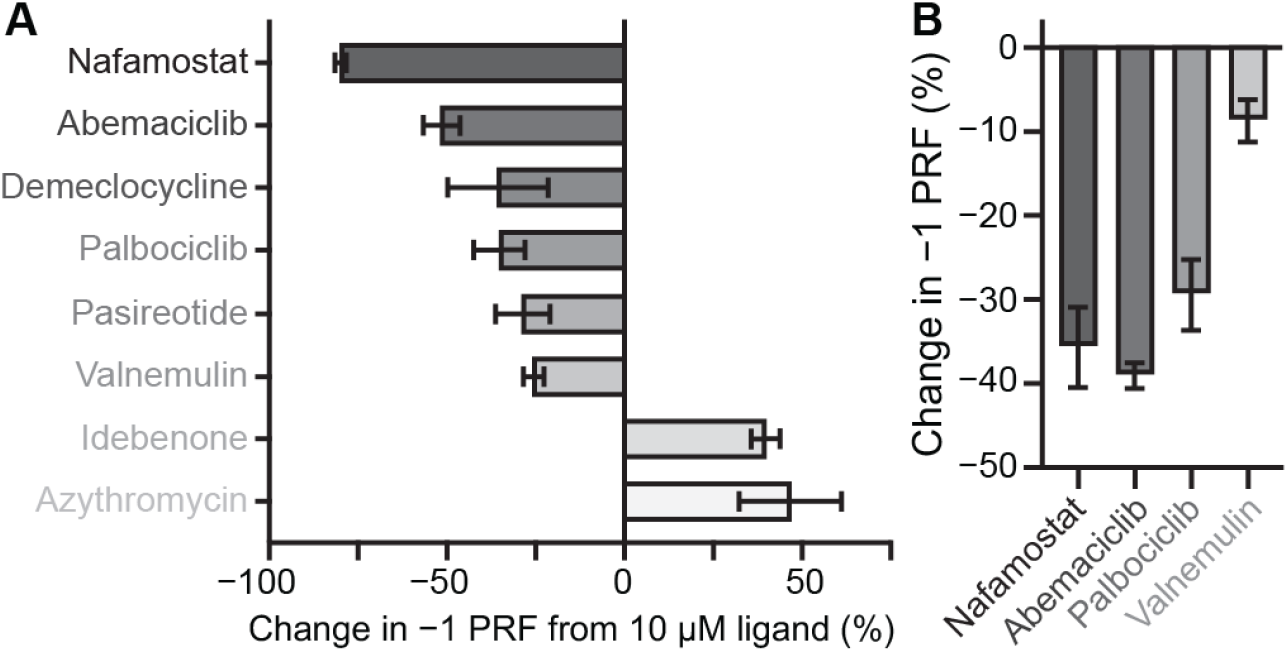
Drugs modulating −1 PRF in SARS-CoV-2 from screening assays. (A) Results from screening a library of 1,814 FDA-approved drugs using a dual-luciferase reporter measured *in vitro* in rabbit-reticulocyte lysate; most of the hits inhibited −1 PRF (dark grey), but some enhanced it (light grey). Error bars represent s.e.m. from 3 replicates. (B) Inhibition of −1 PRF by selected compounds in A549 human lung epithelial cells transfected with a bi-fluorescent reporter system. Error bars represent s.e.m. from 4 replicates.

Nafamostat, a protease inhibitor used as an anticoagulant that also has anti-viral properties and is under clinical investigation for use against COVID-19 (41,42), stood out as the strongest inhibitor. Several of these inhibitors were confirmed to suppress −1 PRF in a cell-based assay, using A549 human lung epithelial cells transfected with a bi-fluorescent frameshifting reporter system (Fig. 3B).

From these screening results, we selected four of the inhibitors to test their effectiveness against different CoV −1 PRF signals in the cell-free assay: nafamostat, the strongest of the inhibitors; abemaciclib and palbociclib, which are CDK4/6 kinase inhibitors approved for use against breast cancer (43,44); and valnemulin, as a representative of an antibiotic approved for veterinary use (45). We also included two compounds previously found to inhibit −1 PRF by the SARS-CoV-2 frameshift signal: MTDB, a compound first identified as an inhibitor for SARS-CoV (24,25) before being shown to be active against SARS-CoV-2 (19,26), and merafloxacin, an experimental fluoroquinolone antibiotic shown to be active against SARS-CoV-2 (11). The structures of each of the compounds used in the tests are shown as insets in Fig. 4. For each compound, the percent change in −1 PRF induced by a final ligand concentration of 20 μM was measured for each frameshift signal in the panel. The specificity to CoV frameshift signals was also assessed by measuring the effects of each compound on −1 PRF stimulated by the frameshift signals from pea enation mosaic virus-1 (PEMV1), which involves a standard 2-stem H-type pseudoknot (46), and HIV-1, which involves a simple hairpin (47,48).

**Figure 4:**
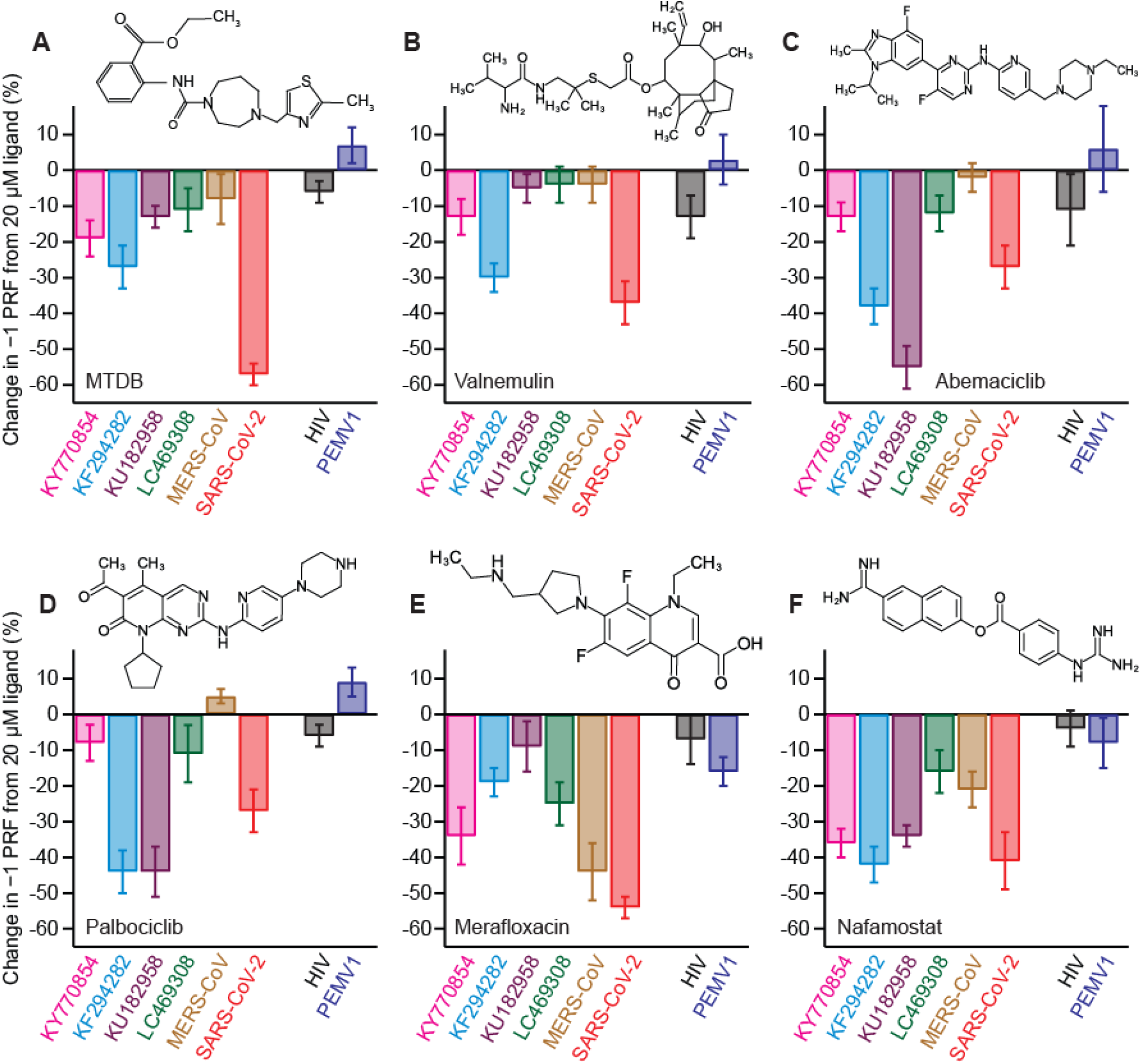
Activity of −1 PRF inhibitors against frameshift signals from different CoVs. (A) Change in −1 PRF efficiency compared to basal levels (Fig. 2) induced by 20 μM MTDB. Remaining panels show the same for (B) valnemulin, (C) abemaciclib, (D) palbociclib, (E) merafloxacin, and (F) nafamostat. In each case, results for CoVs are shown on left, results for specificity controls on right. Experiments performed *in vitro* using dual-luciferase reporter in rabbit reticulocyte lysate. Error bars represent s.e.m. from 3–7 replicates. Insets: chemical structures of inhibitors.

Considering first the results for MTDB (Fig. 4A), we found that it was most effective at inhibiting −1 PRF in SARS-CoV-2 (∼55% decrease); it still induced some modest inhibition for the two alpha-CoVs, KY770854 and KF294282 from clusters 3 and 4 (∼20–30% decrease), but it was ineffective against the beta-CoVs from clusters 1 and 2 (∼10% or less inhibition). The effects of valnemulin (Fig. 4B) followed a similar pattern: most inhibitory for SARS-CoV-2 (∼35% decrease in −1 PRF), modest to minimal inhibition for the two alpha-CoVs (∼15–30% decrease), and no discernable effect on the remaining beta-CoVs. Abemaciclib (Fig. 4C) and palbociclib (Fig. 4D) showed a different pattern of inhibition: they were both most effective against KU182958 from cluster 1 and KF294282 from cluster 4 (∼40–55% decrease in −1 PRF), had a more modest effect against SARS-CoV-2 (∼25–30%), and were minimally effective or ineffective against the remainder. Merafloxacin (Fig. 4E) was most effective against SARS-CoV-2, MERS-CoV, and KY770854 from cluster 3 (∼35–55%), modestly effective against the KF294282 from cluster 4 and LC469308 from cluster 2 (∼20–25%), but ineffective against KU182958 from cluster 1. Finally, nafamostat (Fig. 4F) stood out as having an effect for all the frameshift signals tested: it was most effective at inhibiting −1 PRF in SARS-CoV-2, KU182958 from cluster 1, KY770854 from cluster 3, and KF294282 from cluster 4, leading to similar decreases (∼35– 40%) for all four, and least effective against MERS-CoV and LC469308 from cluster 2 (∼15–20% decrease). For every compound tested, the effects on −1 PRF were minimal (<10%) or consistent with zero for the frameshift signals from HIV and PEMV1 that were used as controls for specificity, with two exceptions that showed a small effect: valnemulin with HIV (13 ± 6% decrease), and merafloxacin with PEMV1 (16 ± 4%).

## DISCUSSION

The results represent, to our knowledge, the first measurements of −1 PRF induced by bat coronavirus frameshift signals. They confirm that there is indeed a programmed frameshift at the expected site between ORF 1a and 1b in each of the viruses chosen to represent the 5 different clusters of bat CoVs. Given that the representative frameshift signals from all clusters except cluster 3 stimulated −1 PRF with efficiency of 25–50%, our measurements suggest that most bat CoVs feature relatively high levels of frameshifting. These values are within the range that has been reported previously for several other coronaviruses (19,30,49,50). The frameshift signal from KY770854 (cluster 3) was a notable outlier, however, stimulating −1 PRF with only 9% efficiency, distinctly below the range typical of CoVs. The origin of this difference is unclear, but it might arise from differences in the pseudoknot structure: the pseudoknot from KY770854 is predicted to contain four stems, rather than the three stems found more commonly in CoV pseudoknots and seen in all the other members of the panel (Fig. S1).

Examining the effects of the inhibitors, our results show that −1 PRF could be suppressed moderately to strongly (>30% decrease) for each of the representative frameshift signals in the testing panel by at least one of the inhibitors: (i) KU182958 from cluster 1 by abemaciclib, palbociclib, and nafamostat; (ii) LC469308 and MERS-CoV from cluster 2 by merafloxacin; (iii) KY770854 from cluster 3 by nafamostat and merafloxacin; (iv) KF294282 from cluster 4 by palbociclib, nafamostat, abemaciclib, and valnemulin; and (v) SARS-CoV-2 from cluster 5 (SARS-like cluster) by MTDB, merafloxacin, nafamostat, and valnemulin. These results provide proof-of-principle that effective small-molecule inhibitors can be found for each of these frameshift signals, and they suggest that the same is likely true for the full range of bat-CoV frameshift signals that these clusters represent.

Several of the inhibitors studied here were moderately to strongly effective against the representatives from more than one CoV cluster: abemaciclib and palbociclib against clusters 1 and 4; valnemulin against clusters 4 and 5; merafloxacin against clusters 2, 3, and 5; and nafamostat against clusters 1, 3, 4, and 5. It was previously noted that merafloxacin was effective at inhibiting −1 PRF in human beta-CoVs but not human alpha-CoVs (11). However, our results show that this pattern does not extend to bat CoVs, as merafloxacin was found to be moderately effective against at least one of the alpha-CoVs, KY770854 from cluster 3. Indeed, there were no obvious patterns in terms of effectiveness against alpha-CoVs versus beta-CoVs for any of the inhibitors active against multiple frameshift signals: in every case, they were effective against at least one alpha-CoV and one beta-CoV, suggesting that the family to which the CoV belongs has little influence on the effectiveness of any of these inhibitors.

The results for nafamostat are especially interesting, because of the breadth and specificity of its effectiveness: it showed inhibition of −1 PRF to some level for all the CoV frameshift signals tested and none of the controls. Furthermore, this inhibition was substantial (>30%) for 4 of the 5 clusters of frameshift signals. These results suggest that nafamostat may be active against a very broad spectrum of CoVs, supporting the notion that targeting −1 PRF holds promise for developing broad-spectrum CoV therapeutics. Intriguingly, nafamostat is in clinical trials for treating COVID-19, putatively because of its activity as a protease inhibitor: it was previously shown to inhibit S-protein mediated membrane fusion and viral infectivity for MERS-CoV (51) and SARS-CoV-2 (41,42). The fact that nafamostat also inhibits −1 PRF may provide it a second, complementary mechanism of action against CoVs. It also suggests there may be utility in pairing nafamostat with nucleotide-analog RdRP inhibitors like remdesivir (52) or molnupiravir (53): suppressing the expression levels of RdRP by inhibiting −1 PRF while also inhibiting the activity of any RdRP that is expressed could lead to significant improvements over the action of either treatment alone.

Finally, we note that the mechanisms of action of the −1 PRF inhibitors we have studied are unclear. Previous work on MTDB found that it bound to the stimulatory pseudoknot from SARS-CoV and reduced its conformational heterogeneity (25), a feature that has been found to be linked to the efficiency with which −1 PRF is stimulated (47,54-57). Indeed, recent work found a predictive correlation between MTDB binding and conformational heterogeneity of the SARS-CoV pseudoknot under tension (58), within the range of forces applied to the mRNA by the ribosome during translation and frameshifting (59,60), suggesting that −1 PRF inhibitors act in part by reducing the conformational heterogeneity of the stimulatory structure. Given the ability of CoV pseudoknots to form multiple folds, some even with different topologies (4,10,20,21,61), it is plausible that inhibitors could stabilize specific folds preferentially, reducing the −1 PRF efficiency. It is also possible that inhibitors may act by modulating interactions between the pseudoknot and the ribosome, given that cryo-EM models of the SARS-CoV-2 pseudoknot complexed with a stalled ribosome identified specific contacts between the two that were found to be important for high −1 PRF efficiency (10). Hopefully results such as those presented here will stimulate more extensive mechanistic studies of −1 PRF inhibitors and their interactions with frameshift signals and ribosomes.

## Acknowledgements

This work was supported by the Canadian Institutes of Health Research (grant reference numbers OV3–170709 and VS1–175534), Alberta Innovates, National Research Council Canada, and National Institutes of Health (grant reference number R01 GM117177). We thank Dr. Niko Grigorieff for providing laboratory space and numerous fruitful discussions.

## Supporting information

**Table S1:**
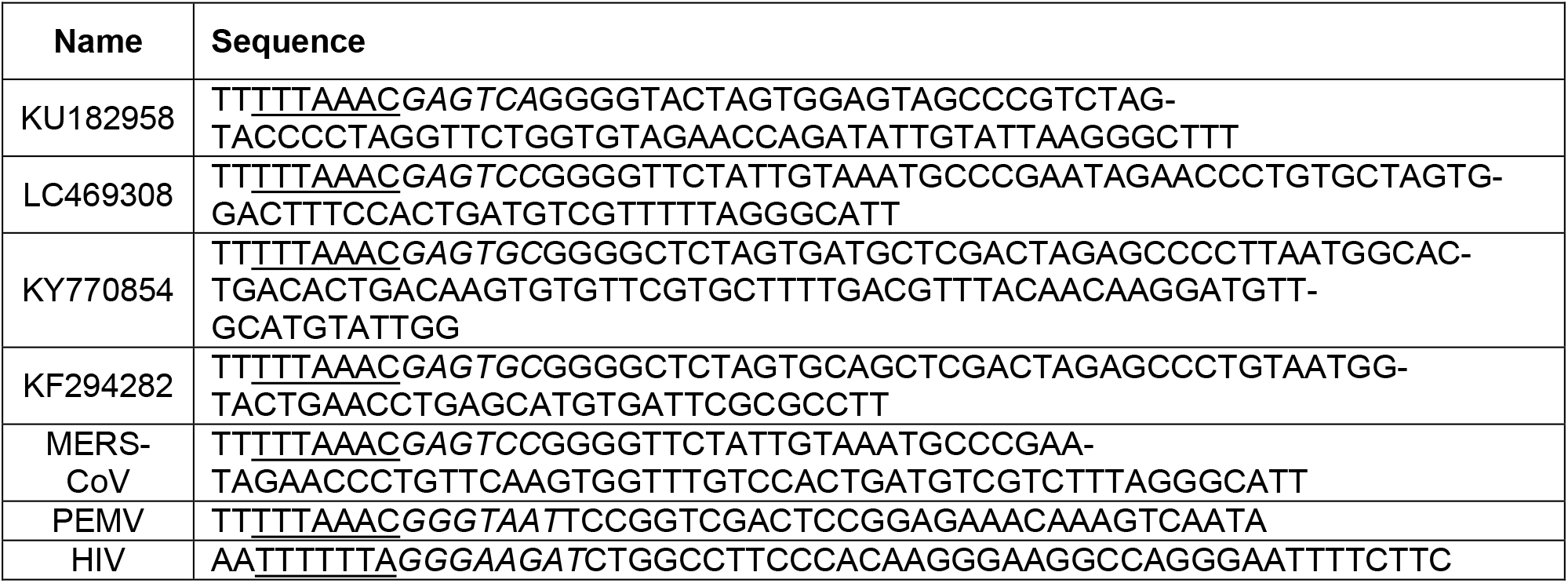
Frameshift signal sequences for panel used in testing inhibitors. Slippery sequence is underlined, spacer is shown in italic font.

**Table S2:**
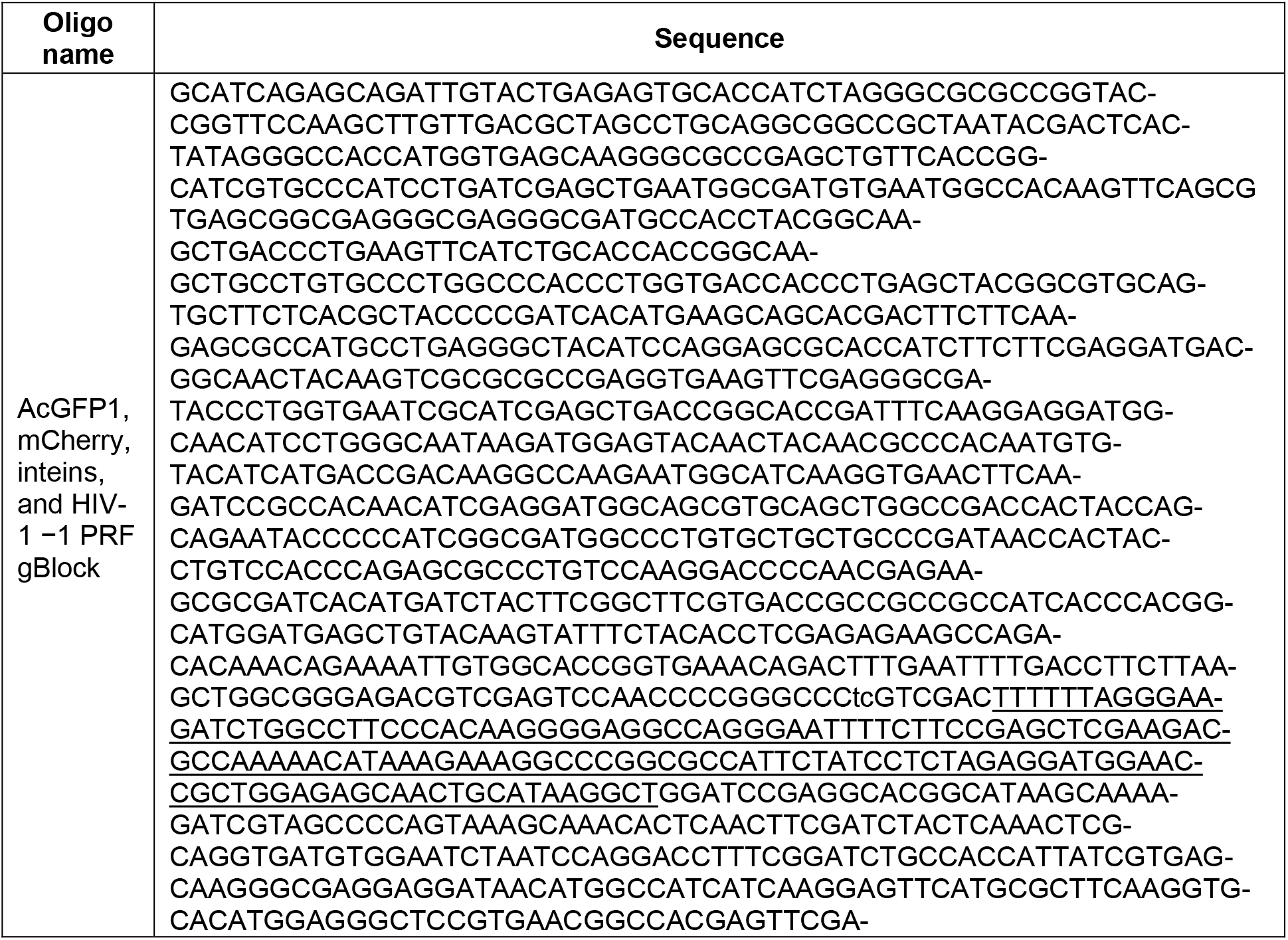
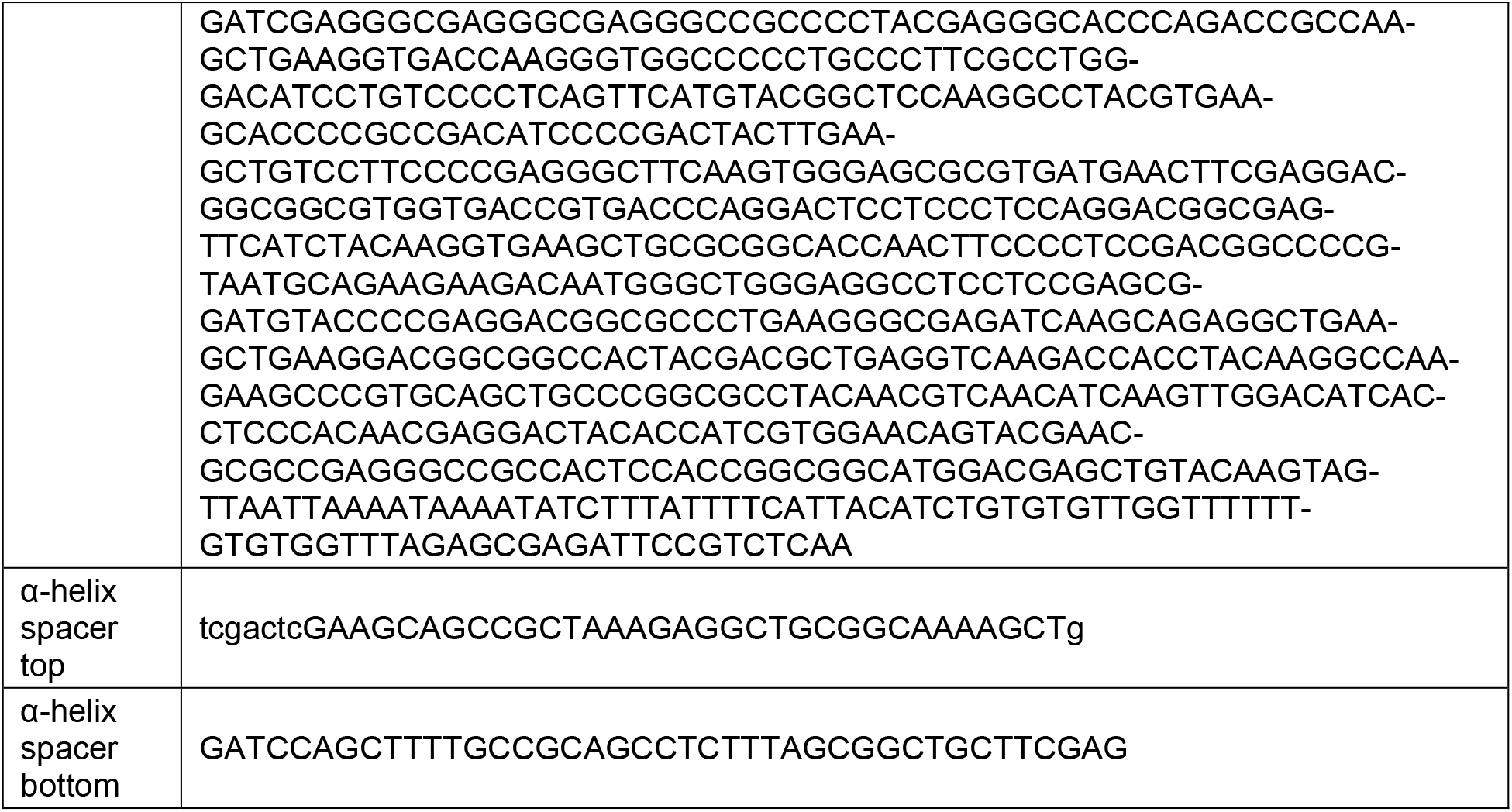
Oligonucleotides for bi-fluorescent reporter construction. Underlined bases in gBlock denote HIV-1 −1 PRF signal, lower-case bases in α-helix spacer denote linkers complementary to restriction sites for cloning.

**Table S3:**
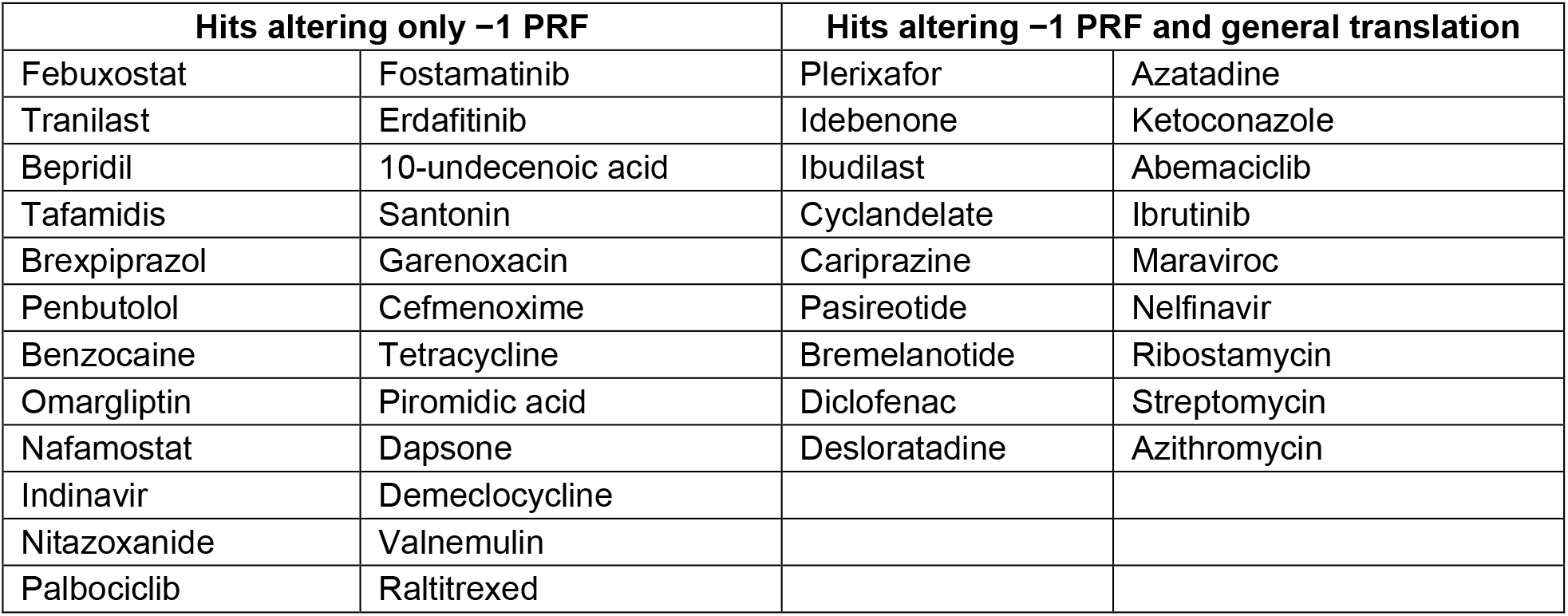
Hits from initial screen of drug library.

**Table S4:**
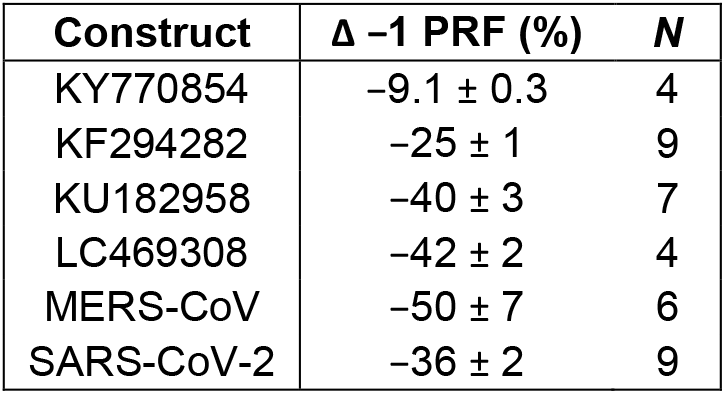
Data in Fig. 2. Error is standard error on the mean for *N* replicates as listed for each construct.

**Table S5:** Data in Fig. 3. *N* = 3 replicates measured in cell-free assays, *N* = 4 in cell-based assays.

| Tests in cell-free assay |  |  | Tests in cell-based assay |  |  |
| --- | --- | --- | --- | --- | --- |
| Compound | $\Delta -1$ PRF (%) | s.e.m. | Compound | $\Delta -1$ PRF (%) | s.e.m. |
| Nafamostat | -80 | 1.4 | Nafamostat | -36 | 5 |
| Abemaciclib | -52 | 5 | Abemaciclib | -39 | 2 |
| Demeclocycline | -36 | 14 | Palbociclib | -29 | 4 |
| Palbociclib | -35 | 7 | Valnemulin | -9 | 3 |
| Pasireotide | -29 | 7 |  |  |  |
| Valnemulin | -26 | 3 |  |  |  |
| Idebenone | +40 | 4 |  |  |  |
| Azithromycin | +47 | 14 |  |  |  |

**Table S5:** Data in Fig. 4. Number of samples *N* listed for each condition.

|  | KY770854 |  | KF294282 |  | KU182958 |  | LC469308 |  |
| --- | --- | --- | --- | --- | --- | --- | --- | --- |
| Compound | $\Delta -1$ PRF (%) | $N$ | $\Delta -1$ PRF (%) | $N$ | $\Delta -1$ PRF (%) | $N$ | $\Delta -1$ PRF (%) | $N$ |
| MTDB | $-19 \pm 5$ | 3 | $-27 \pm 6$ | 5 | $-13 \pm 3$ | 3 | $-11 \pm 6$ | 3 |
| Valnemulin | $-13 \pm 5$ | 3 | $-30 \pm 4$ | 5 | $-5 \pm 4$ | 3 | $-4 \pm 5$ | 3 |
| Abemaciclib | $-13 \pm 4$ | 3 | $-38 \pm 5$ | 4 | $-55 \pm 6$ | 3 | $-12 \pm 5$ | 4 |
| Palbociclib | $-8 \pm 5$ | 3 | $-44 \pm 6$ | 4 | $-44 \pm 7$ | 5 | $-11 \pm 8$ | 4 |
| Merafloxacin | $-34 \pm 8$ | 5 | $-19 \pm 4$ | 5 | $-9 \pm 7$ | 4 | $-25 \pm 6$ | 5 |
| Nafamostat | $-36 \pm 4$ | 3 | $-42 \pm 5$ | 6 | $-34 \pm 3$ | 4 | $-16 \pm 4$ | 3 |

|  | MERS-CoV |  | SARS-CoV-2 |  | HIV |  | PEMV1 |  |
| --- | --- | --- | --- | --- | --- | --- | --- | --- |
| Compound | $\Delta -1$ PRF (%) | $N$ | $\Delta -1$ PRF (%) | $N$ | $\Delta -1$ PRF (%) | $N$ | $\Delta -1$ PRF (%) | $N$ |
| MTDB | $-8 \pm 7$ | 7 | $-57 \pm 2$ | 5 | $-7 \pm 3$ | 4 | $+7 \pm 5$ | 5 |
| Valnemulin | $-4 \pm 5$ | 3 | $-37 \pm 6$ | 5 | $-14 \pm 6$ | 3 | $+3 \pm 7$ | 4 |
| Abemaciclib | $-2 \pm 4$ | 3 | $-27 \pm 6$ | 3 | $-11 \pm 10$ | 6 | $+6 \pm 12$ | 4 |
| Palbociclib | $+5 \pm 2$ | 3 | $-27 \pm 6$ | 4 | $-6 \pm 3$ | 4 | $+9 \pm 4$ | 3 |
| Merafloxacin | $-43 \pm 4$ | 4 | $-54 \pm 3$ | 3 | $-7 \pm 7$ | 5 | $-16 \pm 4$ | 3 |
| Nafamostat | $-21 \pm 5$ | 3 | $-41 \pm 8$ | 4 | $-4 \pm 4$ | 4 | $-8 \pm 7$ | 4 |

**Figure S1:**
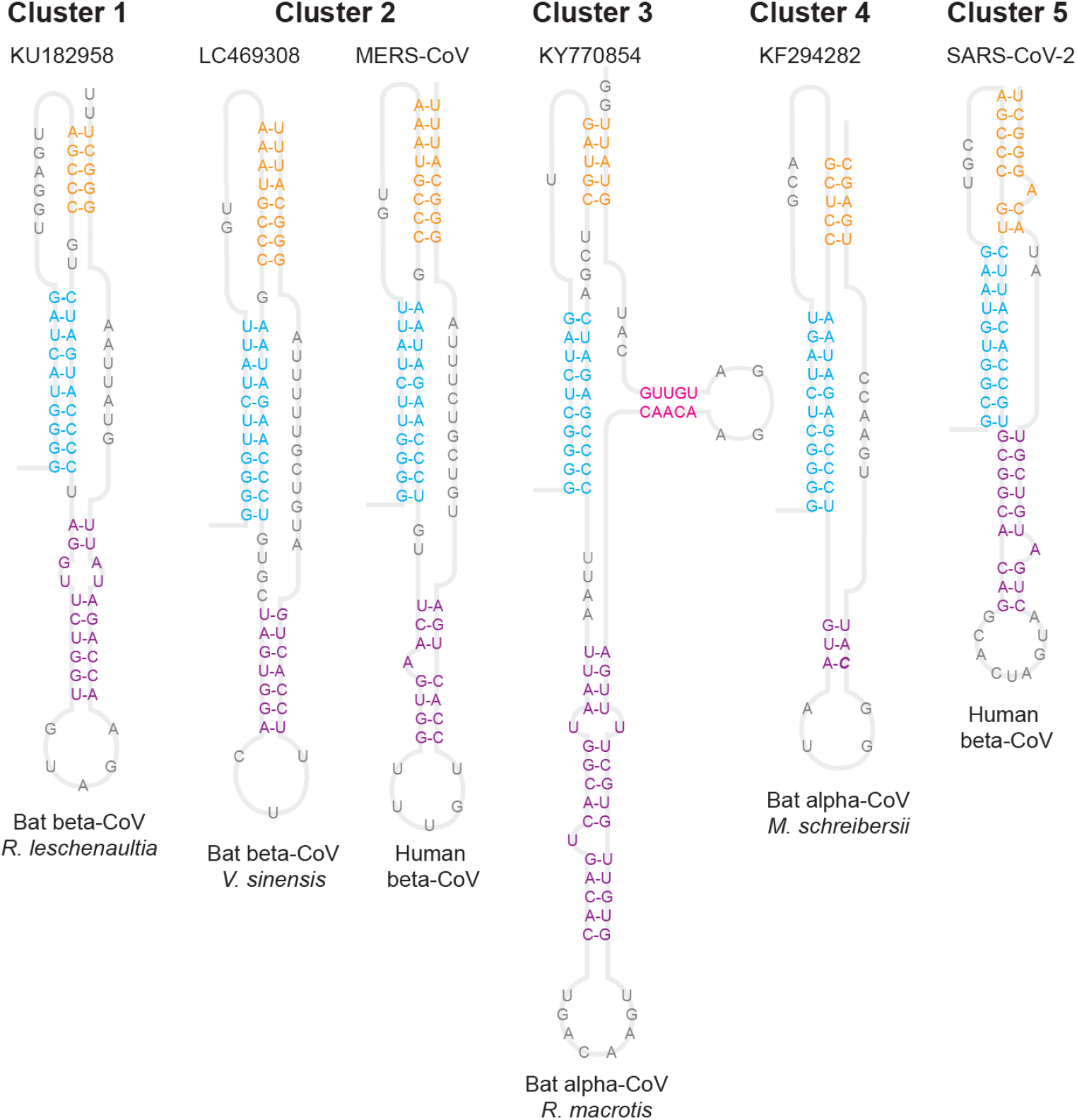
Secondary structure predictions for pseudoknots. Lowest-energy consensus structures homologous to the structures of the SARS-CoV and SARS-CoV-2 pseudoknots. Stem 1: cyan, stem 2: orange, stem 3: purple, stem 4: magenta, loops: grey.

**Figure S2:**
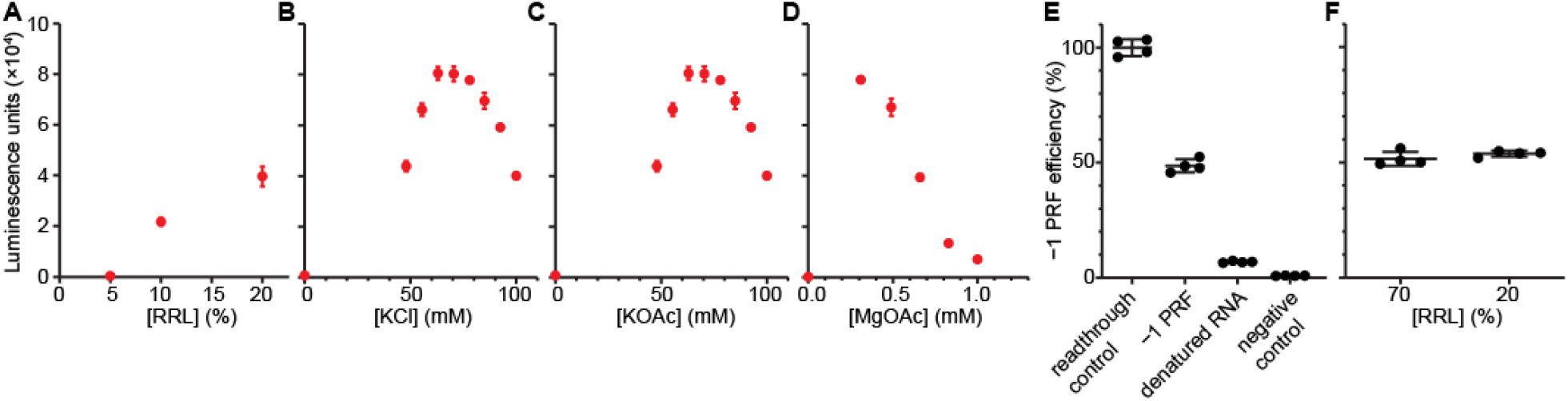
Optimization of dual-luciferase assay for high-throughput screening. We sequentially optimized (A) RRL content, (B) potassium chloride concentration, (C) potassium acetate concentration, and (D) magnesium acetate concentration. Each condition was measured in triplicate. (E) Using optimized conditions, we measured ∼50% −1 PRF efficiency for the SARS-CoV-2 frameshift element. −1 PRF is dependent on the RNA structure, as denaturation of RNA for 5 min at 95 °C followed by snap cooling on ice reduced −1 PRF efficiency nearly to the level of the negative control. (F) Optimized conditions with 20% RRL have the same −1 PRF efficiency as reactions using the manufacturer protocol with 70% RRL (unpaired t-test with Welch’s correction, *p* = 0.26).

